# DNA methylation patterns in breast cancer, paired benign tissue from ipsilateral and contralateral breast, and healthy controls

**DOI:** 10.1101/2025.01.21.634106

**Authors:** Saya R. Dennis, Takahiro Tsukioki, Gannon Cottone, Wanding Zhou, Patricia A. Ganz, Mary E. Sehl, Yuan Luo, Seema A. Khan, Susan Clare

**Affiliations:** Department of Preventive Medicine, Feinberg School of Medicine, Northwestern University, Chicago, IL 60611, USA; Department of Breast and Thyroid Surgery, Okayama University Hospital, Okayama, Japan; Department of Surgery, Feinberg School of Medicine, Northwestern University, Chicago, IL 60611, USA; Center for Computational and Genomic Medicine, The Children’s Hospital of Philadelphia, PA 19104, USA; Department of Pathology and Laboratory Medicine, University of Pennsylvania, Philadelphia, PA 19104, USA; Division of Hematology-Oncology, Department of Medicine, UCLA David Geffen School of Medicine, Los Angeles, CA, United States; Department of Health Policy and Management, Fielding School of Public Health, University of California, Los Angeles, Los Angeles, CA, United States; Department of Computational Medicine, UCLA David Geffen School of Medicine, Los Angeles, CA, United States

**Keywords:** DNA methylation, Breast cancer, Tumor-adjacent normal tissue, Epigenetic profiling, Illumina Methylation EPIC BeadChip, Differential methylation analysis, Hormone-receptor signaling

## Abstract

**Background:** Epigenetic changes, particularly DNA methylation, are crucial to breast cancer development. Tumor-adjacent normal (AN) tissue frequently serves as a reference for characterizing genomic alterations but is reported to share some characteristics with tumors. However, it is unclear whether AN’s epigenetic profiles reflect a predisposition to cancer or a response to the presence of the tumor. We address this gap by systematically comparing methylation profiles of tumor, AN, and matched-benign tissues from both breasts, as well as to healthy donated breast tissue.

**Methods:** We studied four different sample categories from 69 cancer cases: tumor (TU), AN, ipsilateral opposite quadrant (OQ), and contralateral unaffected breast (CUB); and healthy donated breast (HDB) tissue from 182 cancer-unaffected donors. These constitute a “tumor proximity axis” (TPxA): HDB→CUB→OQ→AN→TU. Methylation profiles were assayed using Illumina’s Infinium Methylation EPICv1.0 BeadChip. Differential methylation (DM) analysis was conducted, and the significantly DM CpGs were analyzed for enrichment of transcription factor binding sites (TFBS) and other features.

**Results:** Following data processing and quality control, there were 69 TU, 60 AN, 67 OQ, 68 CUB, and 182 HDB samples for analysis. DM analysis showed distinct methylation profiles of TU relative to benign tissues, whereas case-benign tissues were similar to each other but distinct from HDB. Hypomethylated sites in case-benign versus HDB were enriched for TF binding sites of TP63, GATA3, ESR1, PR, AR, NR3C1, and GREB1. TU hypermethylation events were enriched for Polycomb-repressive complex 2 (PRC2) binding, including EZH2, SUZ12, and JARID2, with hypermethylation enrichment for PRC2-related binding motifs in both ER+ and ER- tumors. TU methylation profiles were otherwise highly distinct by ER status: TFBS enrichment of hypomethylation events for hormone receptor-related pathways in ER+ tumors and for hematopoiesis/immune-related pathways in ER- tumors. We found no differential methylation between benign tissues from patients with ER+ vs. ER- tumors.

**Conclusions:** DNA methylation profiles differ profoundly at two points: tumor to case-benign and case-benign to HDB, with clear distinction between ER+ and ER- tumors. Case-benign tissues are not epigenetically “normal”, are similar across both breasts, and do not differ by ER status of paired tumors.

## Background

Epigenetic changes play a crucial role in cancer development. [1] Among these changes, DNA methylation is one of the most significant due to its impact on gene expression and genome stability. [2–4] DNA methylation occurs at sites in the genome where the sequence is cytosine followed by guanine (CpG), and in which cytosine acquires a methyl group at the 5^th^ position of its ring structure. This modification affects transcription, with hyper-methylation generally suppressing transcription and hypo-methylation promoting it. [2]

In breast cancer research, studying the epigenetic alterations of the tumor requires a non-cancerous reference. The tumor-adjacent normal tissue, typically defined as the morphologically normal tissue located at least 3 cm from the tumor margin [5], is often used as a reference. However, field cancerization––also known as the field effect or field defect––has been a topic of study for decades. Advances in molecular technologies have identified field cancerization in breast cancer, including changes in telomeres, allelic imbalances, DNA methylation, and gene expression, as extensively reviewed by Danforth 2016 [6] and Gadaleta et al. 2022 [7]. These findings support the concept that tumors arise from normal tissue gradually accumulating genomic aberrations, leading to clonal expansion of genomically unstable cells [2, 8]. Earlier DNA methylation studies were limited to a moderate number of CpG loci due to technological constraints [9, 10]; however, with the development of the HumanMethylation 450K (HM450) array, studies examining methylation at over 450,000 CpG loci across the genome became possible. [5, 11–13]. More recently, the EPIC BeadChip array–– surveying nearly twice the CpGs of the HM450––offers a more comprehensive approach.

Breast cancer is heterogeneous, particularly in its subtypes defined by hormone receptor and HER2-neu expression. In previous studies, DNA methylation differences among tumor subtypes have been characterized using the HM450 array [14, 15] but whether the DNA methylation profiles of benign tissues from cancer patients reflect characteristics of their corresponding tumor subtypes remain unexplored. We have employed the more comprehensive EPIC array to fully profile epigenetic differences by estrogen receptor subtype, along with matched benign tissues at various proximities to the tumor and reference control tissues from unaffected donors. We have then systematically compared DNA methylation profiles across tumors, paired benign ipsilateral and contralateral breast samples, and benign breast samples from women with no breast cancer history to investigate epigenetic changes associated with tumor proximity and examine whether case-benign tissues at different locations vary in their similarity or differences from paired tumors. By incorporating both tumor and non-tumor contexts and considering estrogen receptor status, we demonstrate distinct differences in the methylation profile of benign breast tissue remote from the tumor in cases when compared to tissue from healthy controls; we establish that case-benign tissues from three different locations spanning the affected and unaffected control breast have similar methylation profiles; and we identify specific differences in the transition from adjacent normal to tumor tissue based on the ER status of the tumor. Overall, these data provide valuable insights into the complex landscape of breast cancer epigenetics.

## Methods

### Sample processing, library construction and methylation profiling

This study was conducted as a collaboration between investigators at Northwestern University (NU) and the University of California, Los Angeles (UCLA). The case samples (tumor and paired benign) were collected at NU, and the unaffected control samples were obtained by UCLA investigators from the Susan G Komen Tissue Bank at the IU Simon Cancer Center under protocols approved by NU and UCLA’s Institutional Review Boards (IRB # STU00051134 and 16-000853, respectively). Tissues were obtained at NU between September 2011 and September 2022. A total of 334 patients were enrolled, providing mammary tissue samples without impact on clinical care. From cancer-affected participants (cases), tissue was collected from the tumor (TU), the tumor-adjacent normal (AN; at least 2 cm tumor margin), the opposite quadrant (OQ), and from the contralateral unaffected breast (CUB). A total of 79 matched sets were available, with sufficient tissue from the tumor and all three benign tissue locations (hereon referred to as “case-benign” tissue categories).

The mammary tissue samples were stored in Optimal Cutting Temperature compound, which was removed prior to homogenization and DNA extraction using QIAGEN’s AllPrep DNA/RNA Kit. DNA quality was assessed using a NanoDrop 1000 Spectrophotometer; only samples meeting a minimum concentration of 50 ng/μL and a volume of 20 μL were retained for analysis. After excluding samples that did not meet these criteria, DNA from 69 patients was available for methylation profiling using Illumina’s Infinium MethylationEPIC v1.0 BeadChip, covering over 850,000 CpG sites. The assay was performed at the University of Minnesota’s Genomics Center. The first batch assayed included TU, OQ, and CUB samples, and the second batch consisted of AN samples.

To minimize chip-to-chip bias during methylation analysis, we randomized sample placement across the 96-well plates. DNA from different tissues of the same individual was randomly assigned to different columns. To validate that this approach effectively eliminated chip-to-chip bias, we placed randomly selected DNA samples in empty wells and confirmed that methylation profiles were consistent across different chips. This validation confirmed the removal of chip-to-chip bias, enabling us to proceed with the analysis.

Normal mammary tissue was initially collected from 192 cancer-unaffected individuals (controls) who had donated tissue to the Susan G Komen Tissue Bank at the IU Simon Cancer Center. DNA methylation profiling was conducted at UCLA as described in Sehl et al. [16]. To match the age distribution between case and control individuals, the 10 youngest patients were excluded, leaving 182 healthy donated breast (HDB) samples for analysis.

Our five tissue categories form an order along the “tumor proximity axis” (TPxA): HDB → CUB → OQ → AN → TU. (Figure 1a)

**Figure 1.**
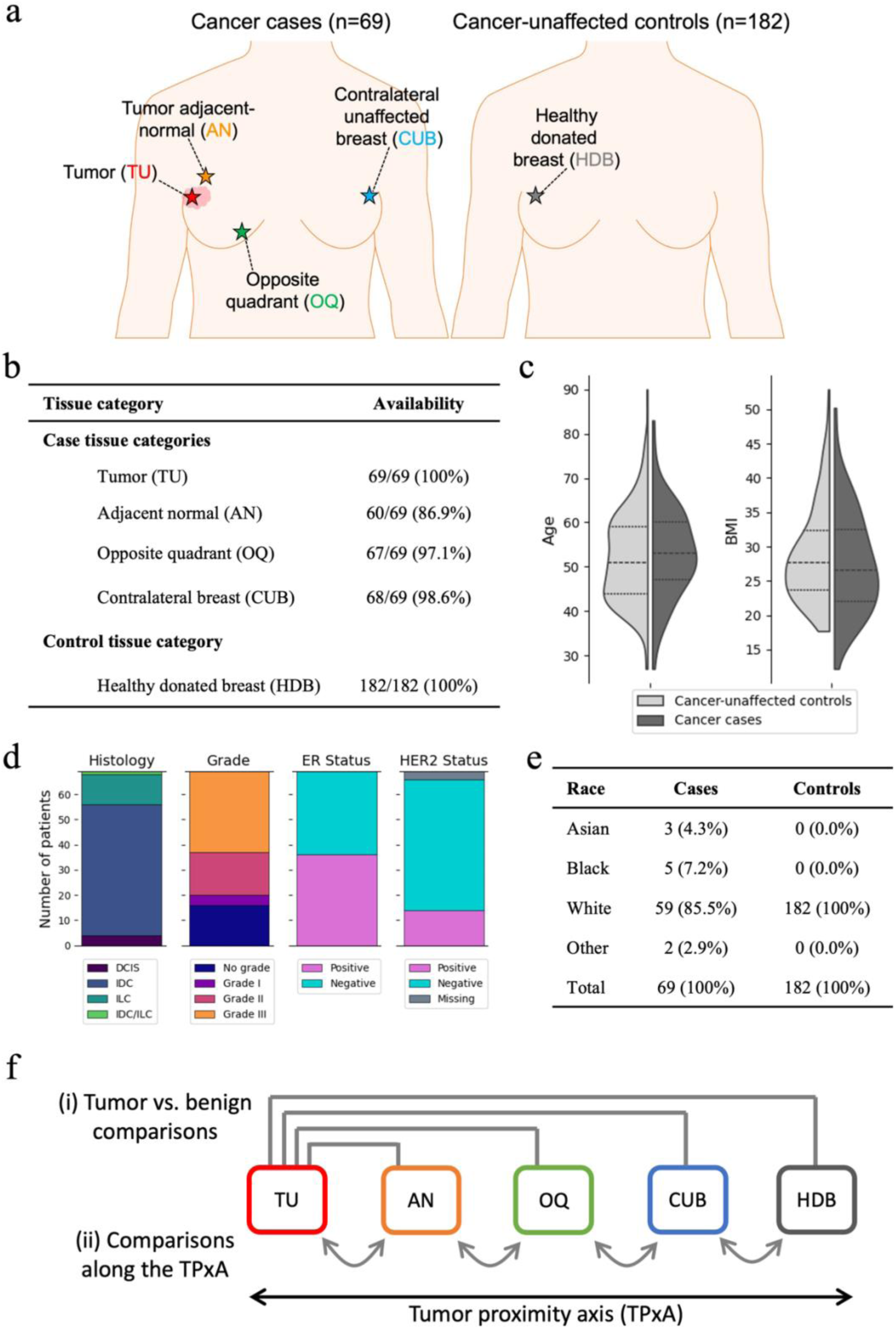
Study design and sample characteristics. Figure 1 legend: **(a)** Illustration of the five tissue categories which were collected. **(b)** Sample availability for each tissue category. **(c)** Age and BMI distributions for cancer cases and cancer-unaffected controls. **(d)** Overview of tumor characteristics for cancer cases. **(e)** Racial distribution of the cohort. **(f)** Illustration of the two sets of four pairwise comparisons: (i) tumor versus the four benign tissue categories, and (ii) comparisons between tissue categories neighboring along the tumor proximity axis (TPxA).

### Data processing and quality control

The raw data was processed using the SeSAMe software (version 1.20.0) [17] with R (version 4.3.0). To minimize technical variation in methylation signal, we applied SeSAMe’s built-in non-linear dye bias correction and used out-of-band signal for background subtraction and detection p-value masking. Samples that had a detection success rate less than 90% were excluded from all following analyses.

Following processing, methylation levels are represented as beta values, which are the ratio of methylated signal intensity to the total signal intensity (methylated plus unmethylated) at each CpG site for each sample. Beta values range from 0 (completely unmethylated) to 1 (completely methylated).

### Differential methylation analysis

Differential methylation between tissue categories was determined by fitting a linear model to the methylation levels, with tissue category, age, and BMI included as covariates. All p-values were adjusted for false discovery rate (FDR) of 0.05 to correct for multiple testing. A stringent significance threshold was applied, defined as the FDR-adjusted p-value < 0.01 and slope < - 0.1 or slope > 0.1, using an F-test for each contrast variable, unless specified otherwise. We conducted two sets of four pairwise comparisons: (1) comparing tumor samples to the four benign tissue categories, and (2) comparing tissue categories that are adjacent along the tumor proximity axis.

### Probe set enrichment analysis

To understand the trends in differentially methylated (DM) CpG sets, we performed probe set enrichment analysis using built-in functions of the SeSAMe software. [17] We queried our DM CpG sets to 5 categories of databases: transcription factor binding sites (TFBS), regions defined in relation to CpG Islands (CGI), histone modification consensus sites (HMconsensus), chromatin states identified by chromHMM [18], and genes (Ensembl). Enrichment was assessed by determining whether the CpGs in our query set were significantly enriched for specific database category probe sets via Fisher’s exact test. Statistical significance was defined as an FDR-adjusted p-value of less than 0.05, and fold-enrichment greater than 1.05.

## Results

### Data quality assessment and patient/sample summary

Prior to quality control of the methylation signals, we started with 69 TU, 60 AN, 69 OQ, 69 CUB, and 182 HDB samples. (Figure 1a) Overall, our samples had uniformly high-quality scores, and only two OQ samples and one CUB sample were removed due to signal detection rates below 0.90. (Figure 1b; Supplementary Figure 1) The cases and cancer-unaffected donors had a mean age of 52.5 (±10.0 SD) and 52.3 (±10.0 SD) respectively, and mean BMI of 28.4 (±7.1 SD) and 28.8 (±6.9 SD), respectively. (Figure 1c) Although age and BMI distributions were comparable between the cancer patients and cancer-free individuals, we correct for these two variables in all differential methylation analyses since these are well-documented confounders. [19, 20] The tumor characteristics of our cases are summarized in Figure 1d, and the racial distribution of all patients is summarized in Figure 1e. The estimated fraction of leukocytes was highest in TU and lowest in HDB. (Supplementary Figure 2)

### The effects of reference selection

Sample clustering via PCA and t-SNE showed that tumor samples were distinctly separated from all case-benign samples (CUB, OQ, and AN), which were in turn highly distinct from HDB samples. (Figure 2a) When examining the number of differentially methylated CpGs (Figure 1f – (i)), we observed more hypomethylation events than hypermethylation events in tumor samples compared to any benign tissue category. (Figure 2b) Additionally, using HDB samples as a reference resulted in nearly double the number of hypomethylation events and moderately fewer hypermethylation events compared to using case-benign samples as a reference. (Figure 2b)

**Figure 2.**
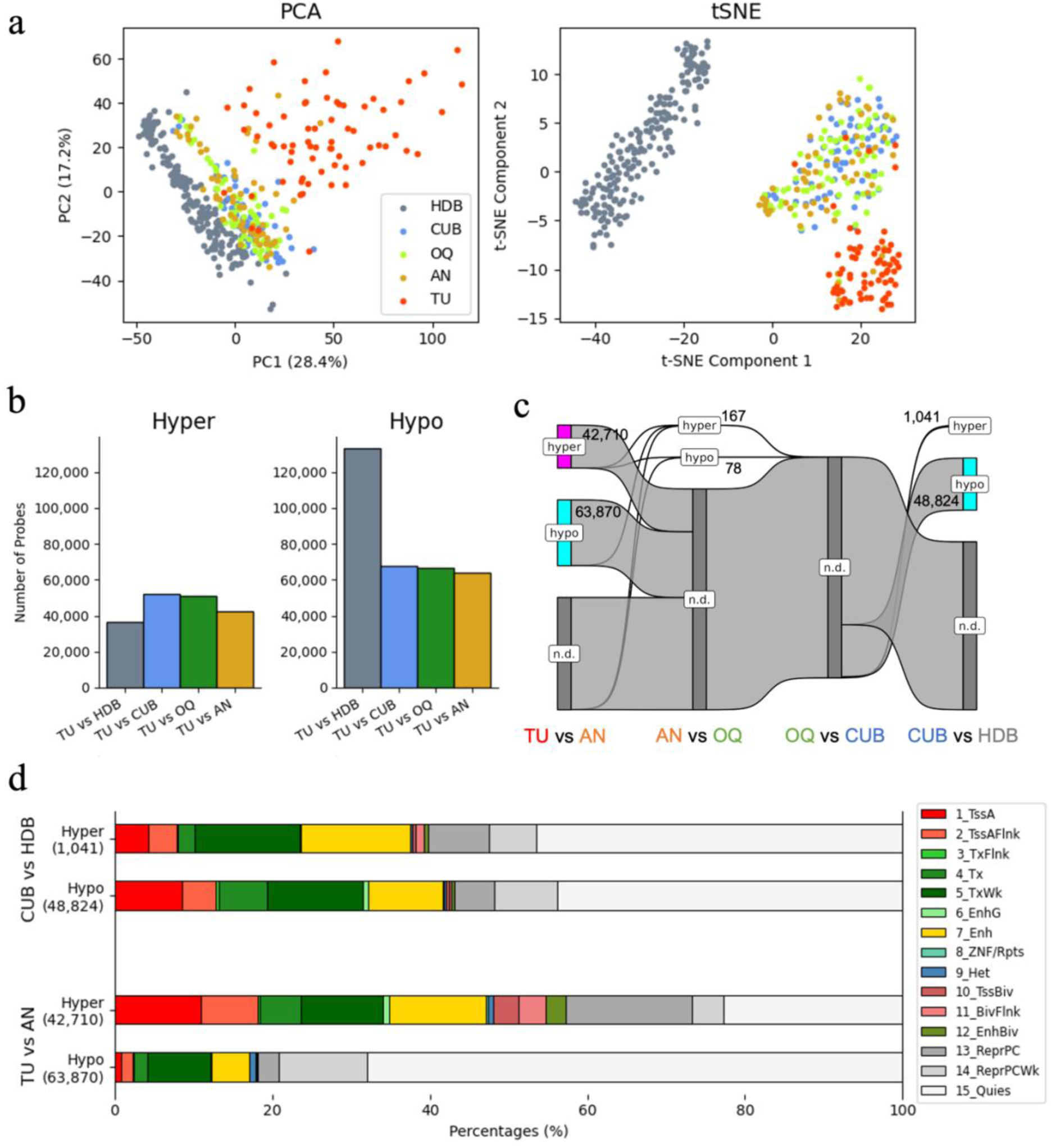
Global methylation trends and number of differentially methylated CpGs. Figure 2 legend: (a) PCA and t-SNE plots of methylation profiles by tissue category. (b) Number of differentially methylated (DM) CpGs in tumor vs. benign tissue comparisons. The second category represents the reference (e.g., in “TU vs CFN,” CFN is the reference category and TU the comparison category). (c) Sankey plot depicting the number of DM CpGs in comparisons along the tumor proximity axis (TPxA). As in panel (b), “A vs B” indicates B as the reference category and A as the comparison category. The height of nodes represents the number of DM CpGs, which is also labeled numerically. The “n.d.” represents CpGs that have no significant difference between categories. (d) Proportions of chromatin states (as defined by chromHMM) for the DM CpGs in the CUB vs HDB and TU vs AN comparisons.

### Differential methylation along the tumor proximity axis

Next, we analyzed the differential methylated (DM) CpGs along the tumor proximity axis (TPxA) (Figure 1f – (ii)). The numbers of hyper- and hypomethylated CpGs are summarized in Figure 2c. TU was distinguished from AN by tens of thousands of both hyper- and hypomethylation events. CUB was also found to be highly distinct from HDB, although this difference was dominated by hypomethylation events. The three case-benign tissue categories (CUB, OQ, and AN) were largely similar to each other, with <300 DM CpGs between AN and OQ and no DM between OQ and CUB (Figure 2c).

The results of the probe set enrichment analysis for DM CpGs along the TPxA are presented in Table 1, with their detailed metrics in Supplementary Tables 1-3. Notably, transcription factor binding site (TFBS) enrichment analysis revealed that hypomethylation events distinguishing CUB from HDB included transcription factor (TF) hits such as TP63 and TP73, along with breast cancer-associated TFs such as ESR1, PR, AR, GREB1, and GATA3. Many of these TFBS enrichment hits were also observed in the hypomethylation events in TU compared to AN, although the enrichment hits were not necessarily driven by the same genes or with the same CpG loci within a gene (Supplementary Figure 3). The pathway enrichment analysis results for genes driving recurrent TFBS enrichment hits in the hypomethylated sites of CUB vs HDB and TU vs AN comparisons are shown in Supplementary Figure 4. Both hyper- and hypo-methylated sets from CUB vs. HDB were enriched for enhancer regions (Table 1). The proportions of hyper- and hypo-methylated CpGs with their corresponding chromatin states defined by chromHMM for the CUB vs HDB and TU vs AN comparisons are shown in Figure 2d.

**Table 1.**
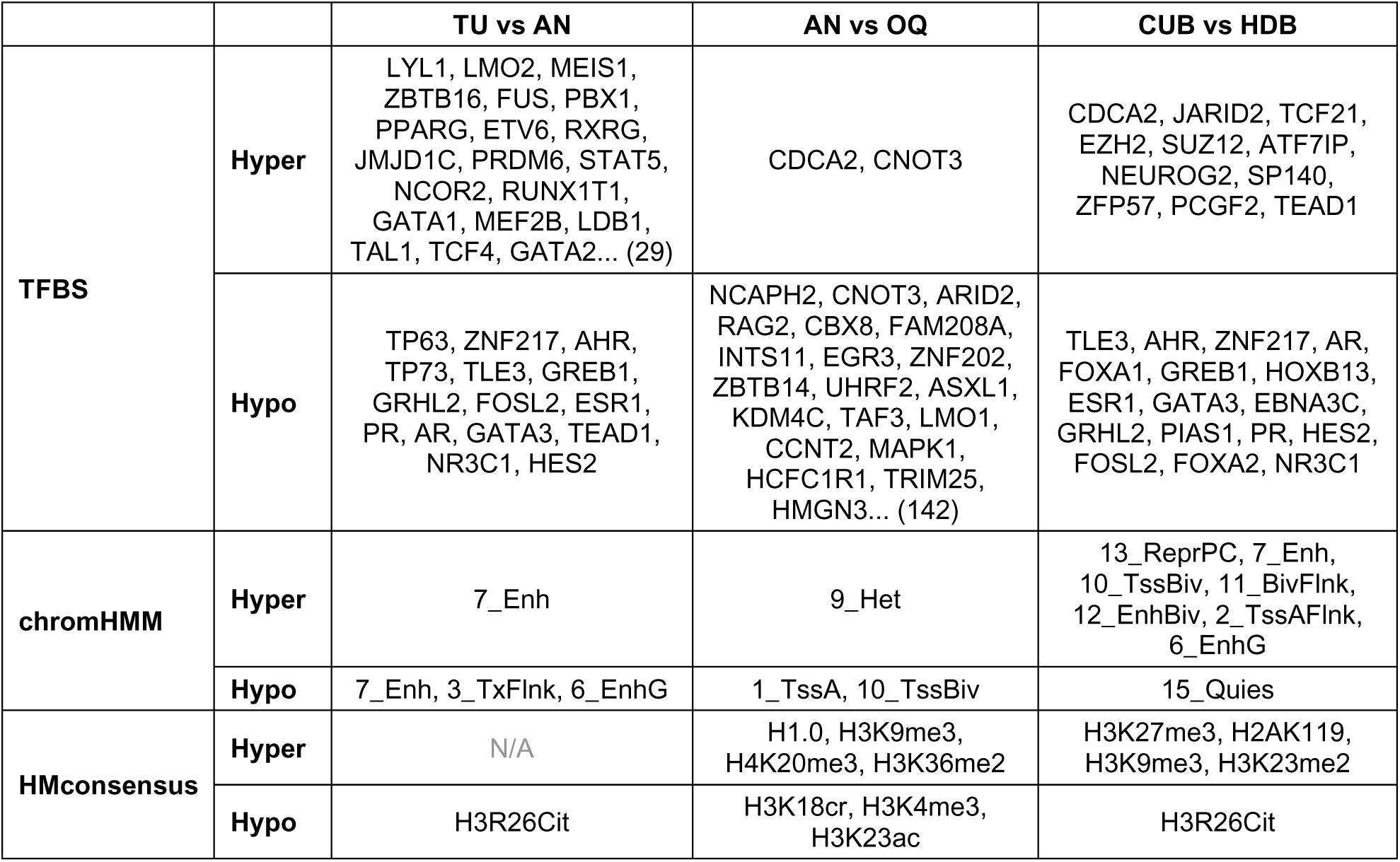
Enrichment analysis results of DM CpG sets along the TPxA.

TFBS enrichment hits in the hypermethylation events in TU compared to AN included EZH2, SUZ12, and JARID2—key components of the Polycomb-repressive complex 2 (PRC2) and particularly the JARID2-associated variant, known for H3K27me3 activity. The hypomethylated CpGs in this comparison were enriched for Quiescent sites (Table 1, Supplementary Table 2).

Although methylation profiles in the case-benign tissue categories were similar to each other, the AN vs OQ comparison yielded 167 hyper- and 78 hypo-methylated CpGs (Figure 2c). The hypermethylation set showed enrichment for binding targets of CDCA2. The hypomethylated CpGs were highly enriched for CpG Islands as well as active transcription start sites (TssA), with 59/78 (75.6%) CpGs mapping to a CpG Island and 64/78 (82.1%) CpGs mapping to active TssA (Table 1, Supplementary Table 2).

Overall, histone modification consensus site (HMconsensus) enrichments in the hypermethylation set corresponded to repressive marks (H1.0, H3K9me3, H4K20me3, H3K27me3, H2AK119, H3K23me2), whereas those in the hypomethylation set corresponded to activation marks (H3R26Cit, H3K18cr, H3K4me3, H3K23ac).

Although most DM CpGs showed a monotonic increase or decrease in methylation levels along the TPxA, a notable subset (6,663 CpGs) was hypomethylated in case-benign tissues compared to both HDB and TU (Supplementary Figure 5: Non-monotonic down & up pattern A). This subset was significantly enriched for genes in the HOXA gene cluster (HOXA2-7; Supplementary Figure 6) and known tumor suppressor genes such as TSC2, EGR1, TP53AIP1, FOXO3 [21–23], RAD51B, and DAXX (Supplementary Table 11). This set was also enriched for consensus sites of H2BK120ub, associated with gene activation, and H4K20me1, associated with both activation and repression (Supplementary Table 11). Additionally, it was significantly enriched for the binding sites of transcription factors TP63, TCF21, TP73, TEAD1, and NEUROG2 (Supplementary Table 11).

### Biomarker-related differences in methylation profiles

PCA and t-SNE plots of case samples (CUB, OQ, AN, TU) revealed that TU samples clustered distinctly by ER status, whereas case-benign samples (CUB, OQ, AN) did not show such separation (Figure 3a). No distinct clustering was observed for HER2 status across any tissue categories (Figure 3b).

**Figure 3.**
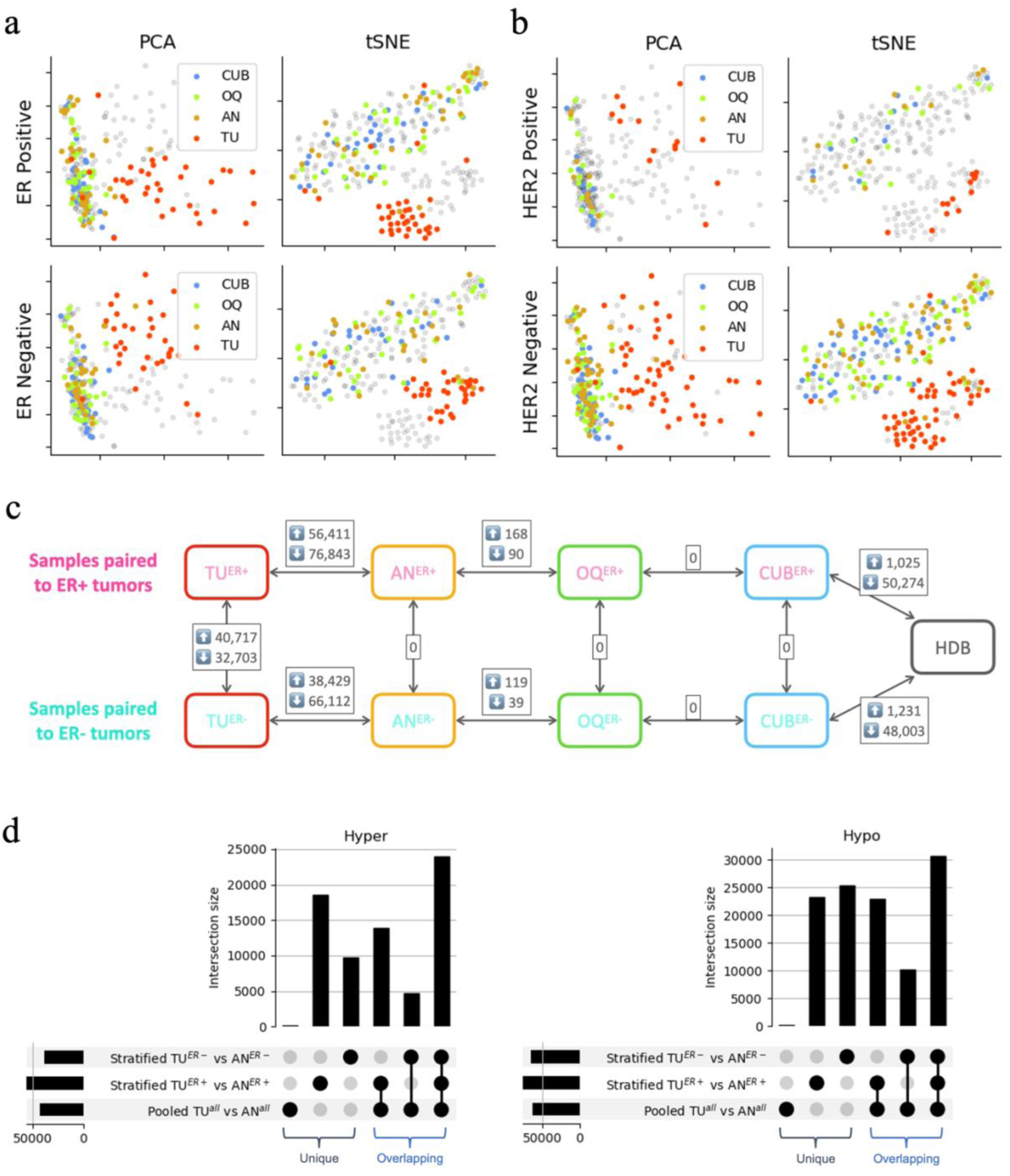
Biomarker-based differences in methylation patterns. Figure 3 legend: **(a, b)** PCA and t-SNE plots of case samples, separated by ER status (a) and HER2 status (b). Light gray points represent samples from the opposing ER or HER2 category, respectively. **(c)** Number of differentially methylated (DM) CpGs in ER-related comparisons. Horizontal comparisons show ER-stratified contrasts along the tumor proximity axis (TPxA), with samples more distal from TU used as the reference. Vertical comparisons represent differences between samples paired to ER+ and ER− tumors. Those paired to ER-negative tumors were used as the reference category. **(d)** Upset plots displaying overlaps between hyper- and hypo-methylated DM CpG sets from pooled or ER-stratified TU vs AN comparison. The vertical bars represent the size of intersections, indicated by the solid circles at the bottom. The horizontal bars in the bottom left indicate the size of each individual DM CpG set by themselves.

Consistent with this lack of clustering, the comparison between HER2+ and HER2-TU samples identified only 5 differentially methylated (DM) CpGs, all hypomethylated in HER2+ samples. These CpGs mapped to genes ERBB2, MIEN1, OSBPL3, EBF1, and PRKN (Supplementary Table 9).

To thoroughly describe methylation trends related to ER status, we defined two sets of contrasts for our differential methylation (DM) analysis: (a) ER-status-based contrasts (vertical contrasts in Figure 3c), and (b) contrasts along the TPxA stratified by ER status (horizontal contrasts in Figure 3c). For ease of notation, we denote the samples that come from patients with ER+ or ER-tumors with superscript such as TU^ER+^ and AN^ER-^. Since unaffected controls have no associated biomarker status, they are simply treated as a single group (HDB).

As for the ER-status-based contrasts, TU^ER+^ vs TU^ER-^ resulted in 40,717 hypermethylated and 32,703 hypomethylated CpGs. No significant DM was detected with case-benign tissues (CUB, OQ, AN) from patients with ER+ versus ER- tumors (vertical contrasts in Figure 3c). The probe-set enrichment analysis results are summarized in Table 2 and Supplementary Tables 6-9. The CpGs hypermethylated in TU^ER+^ compared to TU^ER-^ were enriched for transcription factor binding sites (TFBS) related to chromatin remodeling and epigenetic regulation (HMG20A, ZNF597, NCOR2, JMJD1C), stem cell regulation (LYL1, ETV6, LMO2, RUNX1T1, MEIS1, PBX1, TLX1), and immune modulation (SPIB, NFKB2, RELB, TBX21, BACH2, TERF1). The hypomethylated CpGs were enriched for TFBS associated with hormone receptor (HR) signaling (ESR1, PR, AR, FOXA1, GATA3, NR3C1, HOXB13) and their transcriptional co-regulators (GREB1, PIAS1, TLE3, ZXDC). Both hyper- and hypomethylation events were enriched for enhancer regions identified by chromHMM. The CpGs that were hypomethylated in TU^ER+^ compared to TU^ER-^ were significantly enriched for consensus sites of the histone modification H3R26Cit.

**Table 2.**
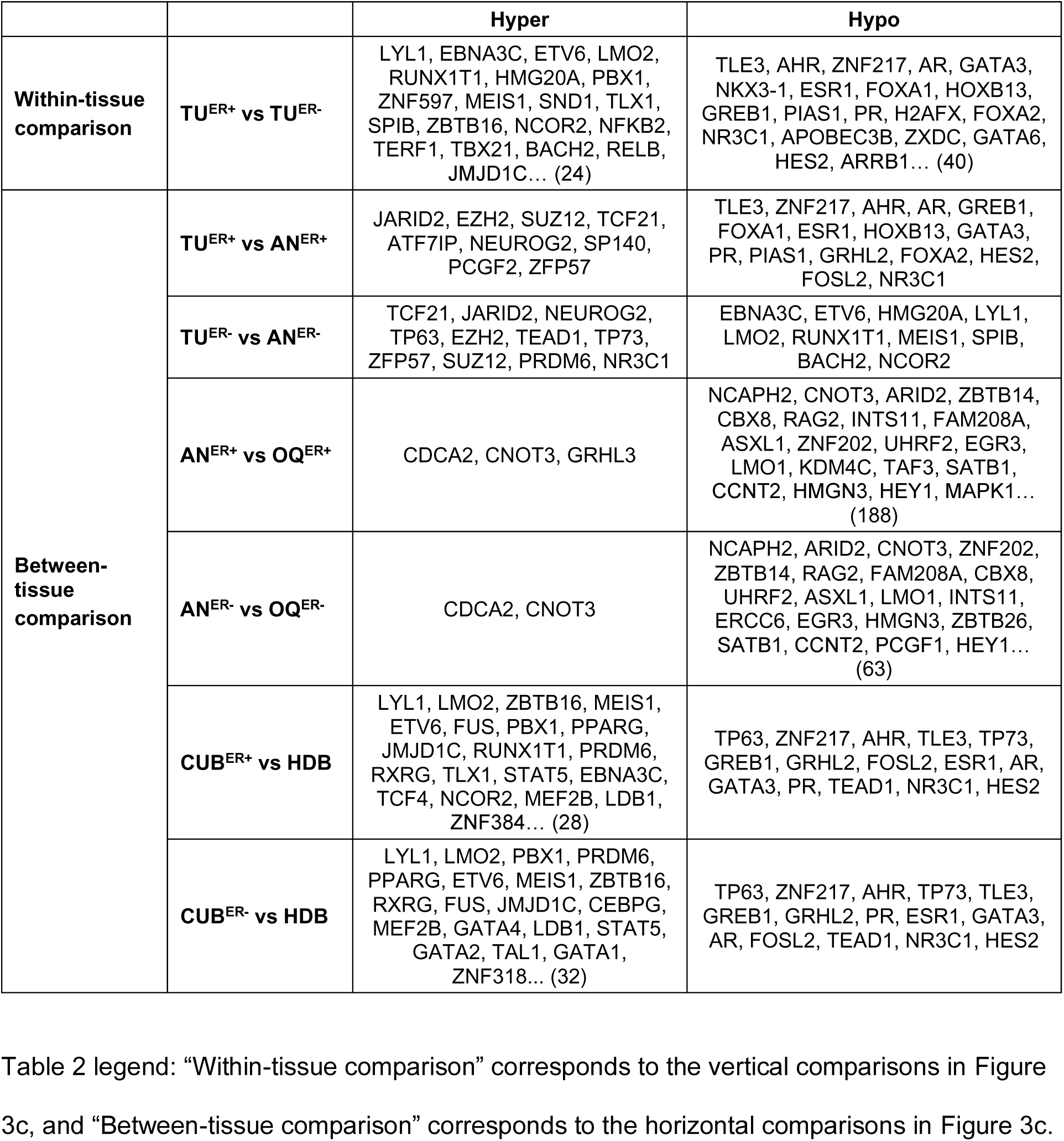
TFBS enrichment analysis results of ER-based comparisons.

Contrasts along the TPxA stratified by ER status (horizontal contrasts in Figure 3c) showed that ER+ contrasts generally yielded a higher number of DM CpGs compared to their corresponding ER- contrasts. Notably, when we assessed the overlaps between ER-stratified and pooled DM CpG sets (Figure 3d), many CpGs were only identified as DM when stratified by ER status (TU^ER+^ vs AN^ER+^ or TU^ER-^ vs AN^ER-^) but not in the combined comparison (Pooled TU^all^ vs AN^all^). (Figure 3d) Specifically, 18,626 hyper- and 23,313 hypomethylated CpGs were unique to the ER+ comparison, while 9,773 hyper- and 25,380 hypomethylated CpGs were unique to the ER- comparison (Figure 3d). Similar trends were seen in the AN vs. OQ comparison, with 46 hyper- and 26 hypomethylated CpGs unique to the ER+ comparison, and 25 hyper- and 7 hypomethylated CpGs unique to the ER- comparison.

TFBS enrichment analysis of TU^ER+^ vs AN^ER+^ and TU^ER-^ vs AN^ER-^ included common hits such as PRC2.2 components (SUZ12, EZH2, JARID2) and other factors (TCF21, NEUROG2). However, there was no overlap between hypomethylation TFBS hit of the TU^ER+^ vs AN^ER+^ and TU^ER-^ vs AN^ER-^ comparisons. Hypomethylated sites in TU^ER+^ vs AN^ER+^ were enriched for HR-related transcription factors (e.g., ESR1, PR, AR, TLE3, ZNF217, GREB1, FOXA1, HOXB13, GATA3). Conversely, hypomethylated sites in TU^ER-^ vs AN^ER-^ were enriched for TFBS of transcriptional regulators (e.g., HMG20A, NCOR2, RUNX1T1) and stem cell regulators (e.g., LYL1, ETV6, LMO2, MEIS1). Histone modification consensus site enrichment for H3R26Cit was only observed in TU^ER+^ vs AN^ER+^ and not in TU^ER-^ vs AN^ER-^.

Interestingly, although there was no significant DM between benign tissues from patients with ER+ and ER- tumors (AN^ER+^ vs AN^ER-^; OQ^ER+^ vs OQ^ER-^; CUB^ER+^ vs CUB^ER-^), the number of DM CpGs differed between the CUB^ER+^ vs HDB comparison and the CUB^ER-^ vs HDB comparison (Figure 3c). When we further investigated any trends in the non-overlapping hyper- or hypo-methylated CpGs between these comparisons (Supplementary Figure 6), we found that the CpGs that were only hypomethylated in the ER+ comparison were enriched for TFBS of FOSL2, TLE3, TEAD1, NR3C1, and AR, whereas those only hypomethylated in the ER-comparison were enriched for TFBS of TP63, GREB1, AHR, TP73, PR, and GATA3 (Supplementary Table 11).

## Discussion

This study provides the most comprehensive characterization to date of methylation profiles in tumors, three categories of case-benign tissues from 69 women, and normal breast tissue from 182 controls unaffected by cancer. Using the EPICv1 BeadChip technology, we assayed over 850,000 CpG sites across the genome.

Our findings reveal significantly different DNA methylation profiles at two critical points along the tumor proximity axis (TPxA): between healthy donated breast (HDB) and contralateral unaffected breast (CUB) tissues, and between adjacent-normal (AN) and tumor (TU) tissues. This difference was evident in PCA/t-SNE visualizations and the count of differentially methylated (DM) CpGs. (Figure 2a, c) Notably, TFBS enrichment of CUB-HDB differences underscored important breast cancer-related factors, including ESR1, PR, AR, GREB1, and GATA3. (Table 1) These results indicate the unaffected tissues in cancer cases are not epigenetically “normal”. In all our DM comparisons, we generally observed a larger number of hypomethylation events than hypermethylation events in TU compared to other tissue categories. This finding aligns with our current understanding of DNA methylation in tumors, which is characterized by global hypomethylation and localized hypermethylation. [24, 25]

Theoretically, two possible explanations could account for the observed differences between HDB and case-benign tissues. The first, more common explanation is that case-benign tissues reflect the conditions under which tumors develop. Tumor initiation is rarely a single event; instead, it results from the accumulation of multiple factors, both exogenous (e.g., behavioral, environmental) and endogenous (e.g., aging, genetic susceptibility, inflammation) [26]. These factors may predispose larger regions of tissue (or the whole body) to malignancy, leading to epigenetic patterns associated with cancer risk [27]. The second explanation suggests reversed causality, where the epigenetic profiles in case-benign tissues develop in response to the tumor’s presence. Such tumor-responsive trends have primarily been studied in the context of tumor microenvironment [28]. The fact that the methylation profiles of AN, OQ and CUB are remarkably similar favors the first explanation. A broad response to the tumor cannot be ruled out, but is considered unlikely especially in the contralateral breast.

Although further research is required to clarify the mechanisms behind our findings, we provide novel data supporting a critical concept: while the TU vs AN comparison remains informative for certain questions, they do not fully capture epigenetic distinctions between cancerous and normal tissues. Given the important role of the epigenetic environment on gene expression and protein content, these data have similar implications for molecular profiling in these additional dimensions.

The hypermethylation events distinguishing TU from AN were enriched for PRC2 transcription factor binding sites (TFBS), as well as chromHMM’s Polycomb-repressed sites, reinforcing PRC2’s known role in breast and other cancers. [29–31] Notably, we observed significant TFBS enrichment not only for core PRC2 components, EZH2 and SUZ12, but also of its optional subunit, JARID2, which recent studies have shown to promote breast tumorigenesis [32–35]. A study in prostate cancer suggests that PRC2 and DNA methylation may serve as a switch between transient and permanent gene silencing [30]. This may explain why PRC2 targets show unique enrichment in TU-AN differences but not in CUB-HDB differences. Certain genes transiently silenced by PRC2 in HDB, CUB, OQ, and AN may become locked in a repressed state via DNA methylation in TU. Although no specific PRC2 variant has been linked directly to DNA methylation, our findings imply the JARID2-associated variant (PRC2.2).

Interestingly, using HDB as a reference to TU resulted in 15-30% fewer hyper-methylated CpGs than when case-benign tissues were used as a reference (Figure 2b). This reduction is due to a subset of CpGs being hypomethylated in case-benign tissues compared to both HDB and TU (“Non-monotonic down & up pattern A” in Supplementary Figure 5). One notable group showing this pattern was the HOXA gene cluster (Supplementary Figure 7). HOXA genes have been demonstrated to be hypermethylated in breast tumors [36], with some showing tumor-suppressor effects [37, 38], which may explain their hypomethylation in case-benign tissues to counteract the proliferative signals. Some CpGs in this set also mapped to other tumor-suppressor genes, including BAP1 [39], FBXW7 [40], FOXO3 [21–23], and TSC2 [41].

The TU^ER+^ vs TU^ER-^ comparison exhibited striking methylation profile differences, with tens of thousands of DM CpGs, while TU^HER2+^ vs TU^HER2-^ showed only five DM CpGs (Figure 3a-c). This aligns with previous studies that identified fewer DM CpGs in HER2+ tumors compared to luminal subtypes [42, 43], although the exact CpG counts vary due to assay differences and threshold choices. Interestingly, no differential methylation was observed between benign tissues from women with ER+ vs. ER- tumors, suggesting that differences related to ER status of the tumor emerge later in the course of tumor development. While prior studies reported that gene expression in cancer-adjacent tissue can reflect tumor subtype [44], we observed minimal epigenetic differences. Hypomethylated CpGs in TU^ER+^ vs TU^ER-^ were, as expected, enriched for hormone receptor (HR)-related TFBS (Table 2).

Importantly, stratifying the TU-AN comparison by the tumor’s ER status revealed tens of thousands of DM CpGs that were not detected when ER groups were combined (Figure 3d). By comparing the stratified enrichment results (Table 2) with the pooled results (Table 1), we identified which findings were consistent across ER groups and which were distinct. For example, PRC2.2 TFBS, quiescent sites, and H3K27me3/H2AK119 sites were consistently enriched in both TU^ER+^ vs AN^ER+^ and TU^ER-^ vs AN^ER-^ comparisons. In contrast, enrichment of H3R26Cit sites and many hypomethylation TFBS hits were predominantly seen in ER+ comparisons. These findings highlight the importance of stratifying by ER status to more accurately capture the biological diversity among breast tumors.

The hypomethylated CpGs in the CUB-HDB and TU-AN comparisons shared many enrichment hits, such as TFBS of HR signaling proteins and cofactors (e.g., ESR1, PR, AR, GREB1, GATA3, ZNF217, TLE3, GRHL2) as well as the consensus histone modification sites of H3R26Cit (Table 1). These findings gain added value when interpreted in conjunction with those of the tumor ER status-related comparisons. Specifically, many enrichment hits in the TU-AN comparison appear specific to ER+ samples: HR-related TFBS and H3R26Cit sites are enriched in TU^ER+^ vs TU^ER-^ and TU^ER+^ vs AN^ER+^, but not in TU^ER-^ vs AN^ER-^ (Table 2; Supplementary Tables 6-9). However, in the CUB vs HDB comparison, HR-related TFBS and H3R26Cit enrichment are present in both CUB^ER+^ vs HDB^all^ and CUB^ER-^ vs HDB^all^ comparisons. This suggests that case-benign tissues are experiencing elevated HR-related signaling, regardless of the ER status of the tumor.

Our exploration of the non-overlapping DM CpGs between the CUB^ER+^ vs HDB and CUB^ER-^ vs HDB comparisons showed that the non-overlapping DM CpGs between these two comparisons were enriched for many HR-related factors, such as AR, TLE3, GATA3, GREB1, PR, and TP63 (Supplementary Table 11). These findings imply that, although there were no significant differences in the direct DM analysis between CUB^ER+^ vs CUB^ER-^, there may be subtle signs of the paired tumor’s ER status in the case-benign epigenetic profiles as well.

The enrichment of H3R26 citrullination (H3R26Cit) sites highlights the need for further research into potential interactions between DNA methylation and H3R26Cit in breast cancer. H3R26Cit is induced by estradiol [45] and facilitates ER binding to DNA [46]. While several studies have examined links between DNA methylation and histone methylation/deacetylation [47–49], no direct interactions between DNA methylation and histone citrullination have been reported. Our observed enrichment of H3R26Cit sites in both CUB-HDB and ER-based TU-AN comparisons, together with these known links between estrogen pathways and H3R26Cit, suggest a promising direction of further research.

One limitation of our study is that the majority of our patient population is of European descent, which restricts the generalizability of our findings across other racial and ethnic groups. Another concern is potential batch effect, as HDB tissue processing and methylation assays were conducted at a different institution. We mitigated this by applying probe quality masking, dye bias correction, detection p-value masking, and background subtraction. [17, 50] Furthermore, breast cancer-related trends in our TFBS enrichment results suggest the major findings reflect meaningful biology, though technical artifacts in a small subset of DM CpGs, particularly in HDB comparisons, cannot be fully excluded. Finally, our study lacked external validation. Future investigations would benefit from either experimental validation (e.g. RNA expression analysis) or the validation of our findings in an independent cohort.

## Conclusions

Our study revealed substantial epigenetic differences at two steps along the tumor proximity axis, underscoring the importance of tissue context and ER status in breast cancer biology. By comparing both case-benign and unaffected control tissues, we demonstrate that case-benign tissues display hormone receptor-related epigenetic changes independent of the tumor’s ER status, suggesting a predisposition to malignancy and breast tissue remote from the tumor. Additionally, our findings of PRC2 and H3R26 citrullination enrichment, particularly in ER+ tumors, hint at a complex interplay between DNA methylation and histone modifications, providing a basis for future studies to explore these mechanisms in breast cancer progression. Together, these insights provide valuable directions for understanding the multifaceted roles of epigenetic regulation in breast tumorigenesis and underscore the importance of reference selection and ER context in capturing the full diversity of breast cancer biology.

## Supplementary Figures

**Supplementary Figure 1.**
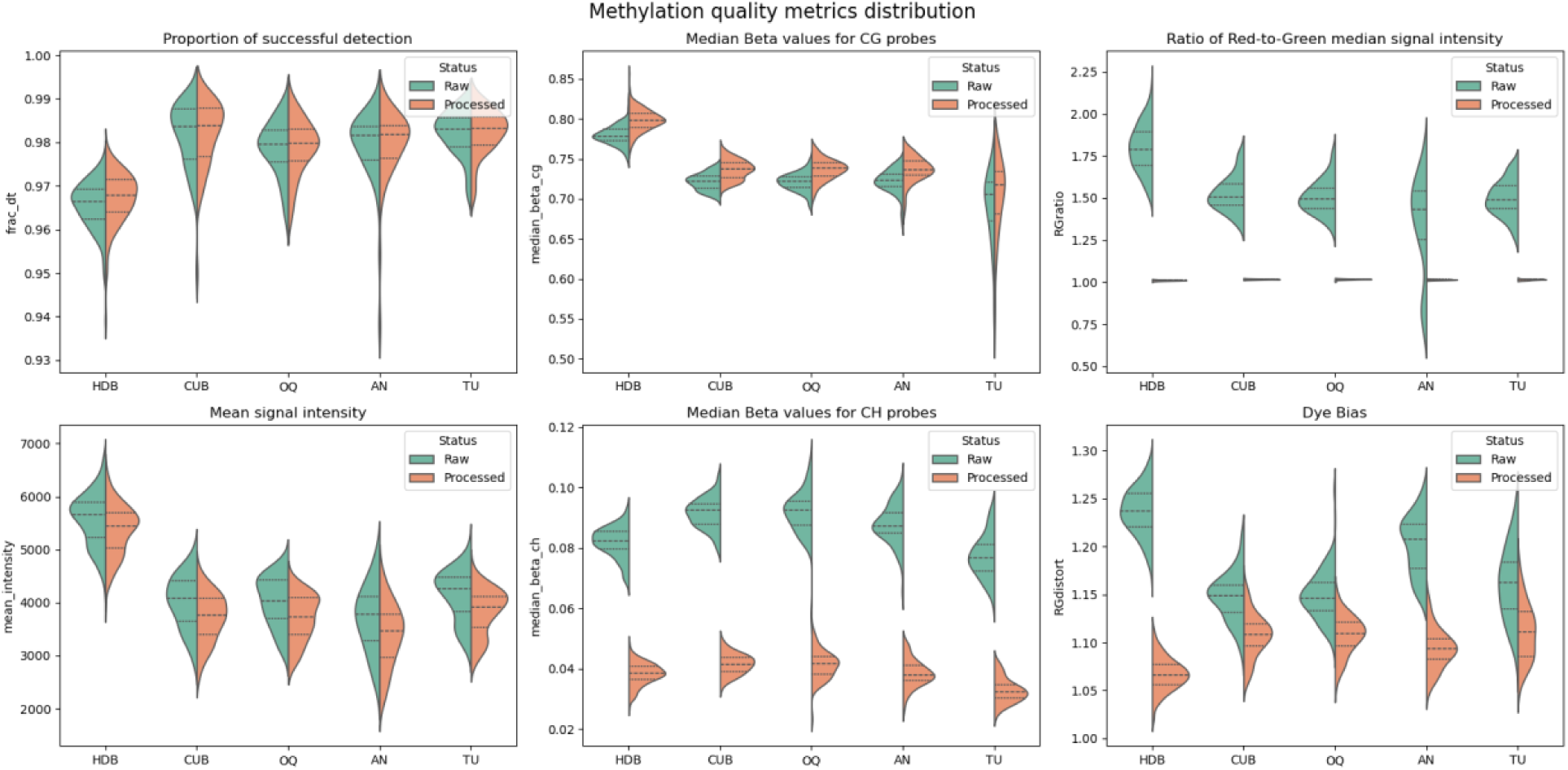
Distributions of data quality metrics pre- and post-processing.

**Supplementary Figure 2.**
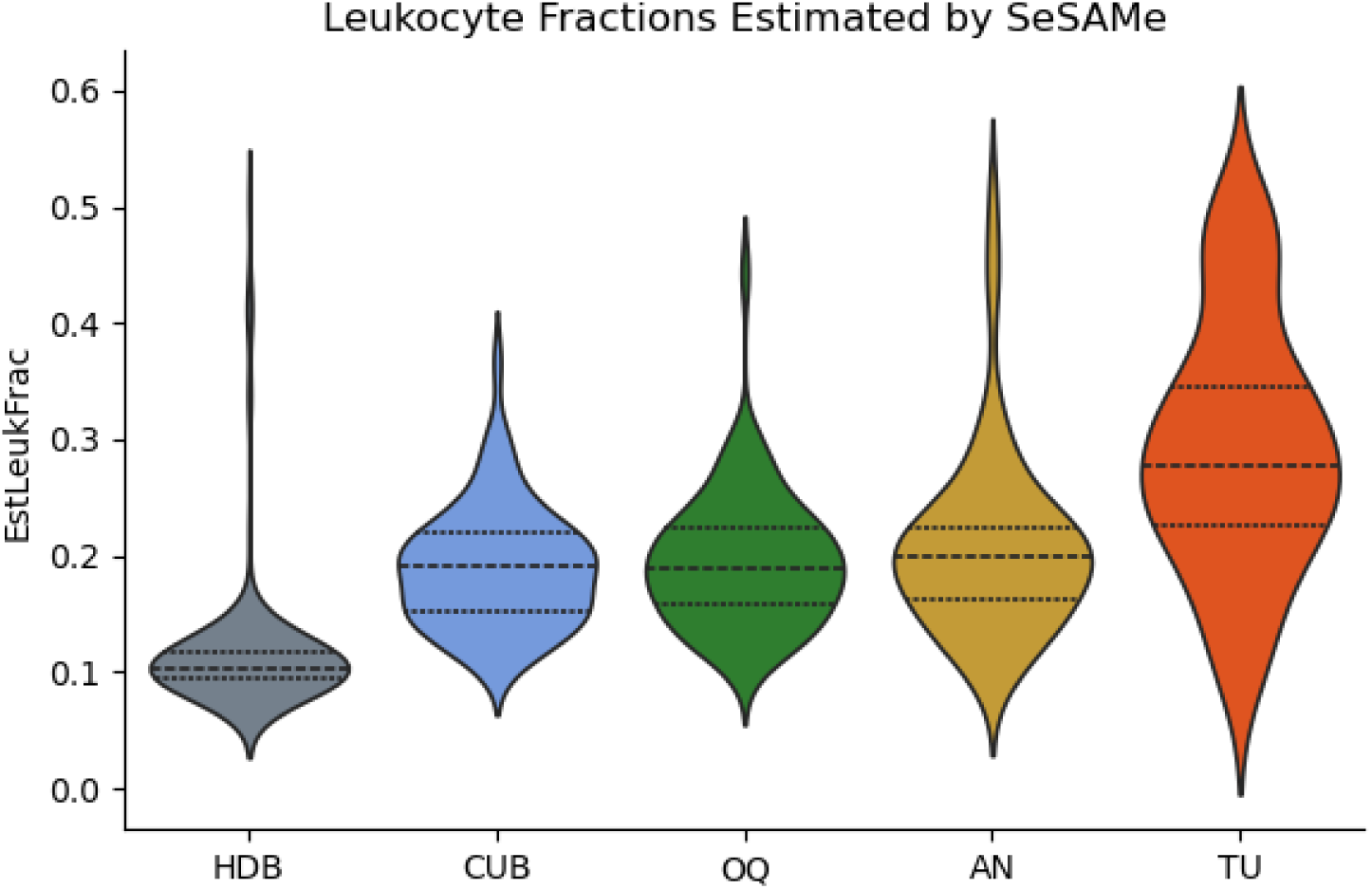
Estimated fraction of leukocytes.

**Supplementary Figure 3.**
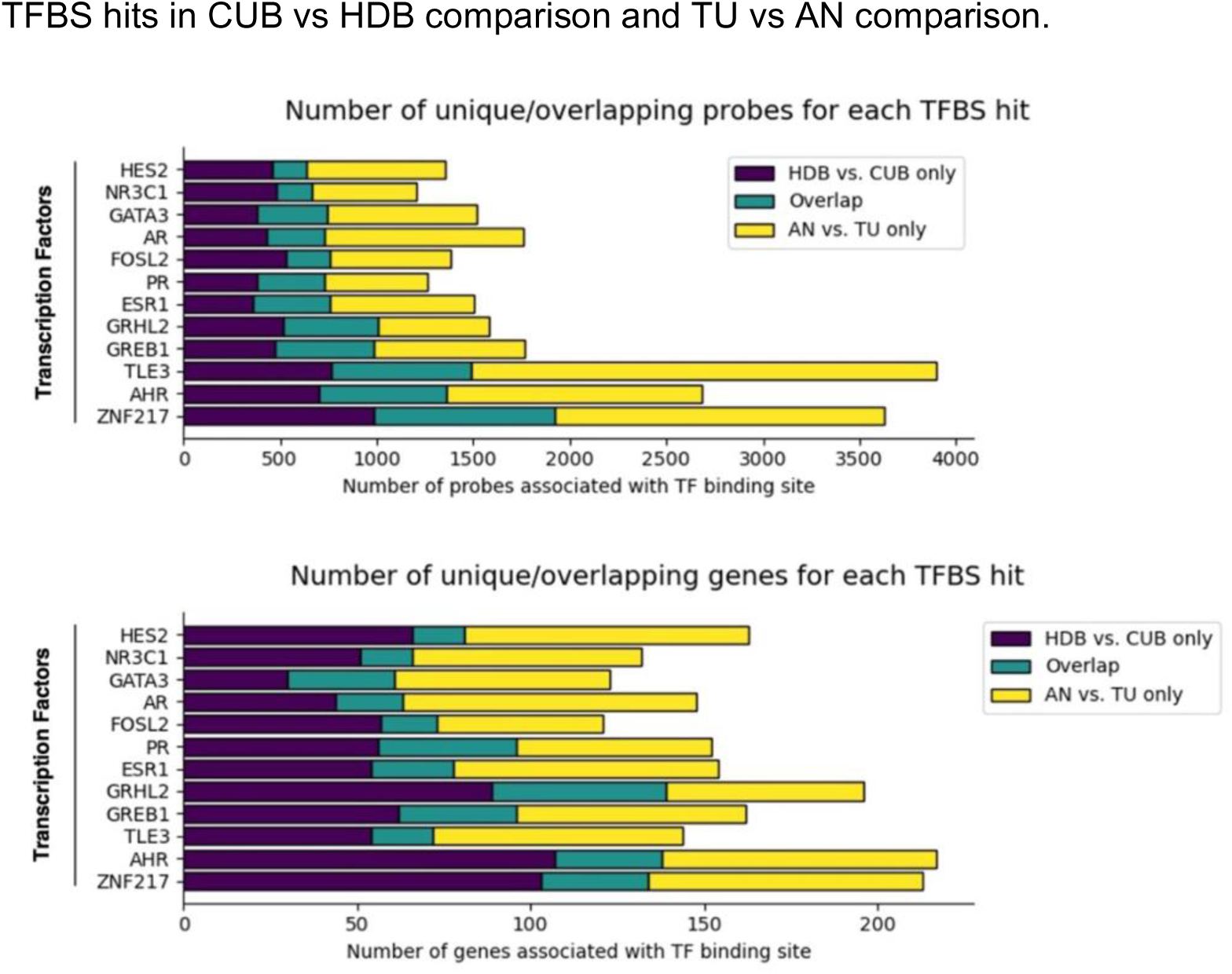
Overlaps in probes and genes driving recurrent hypomethylation TFBS hits in CUB vs HDB comparison and TU vs AN comparison.

**Supplementary Figure 4.**
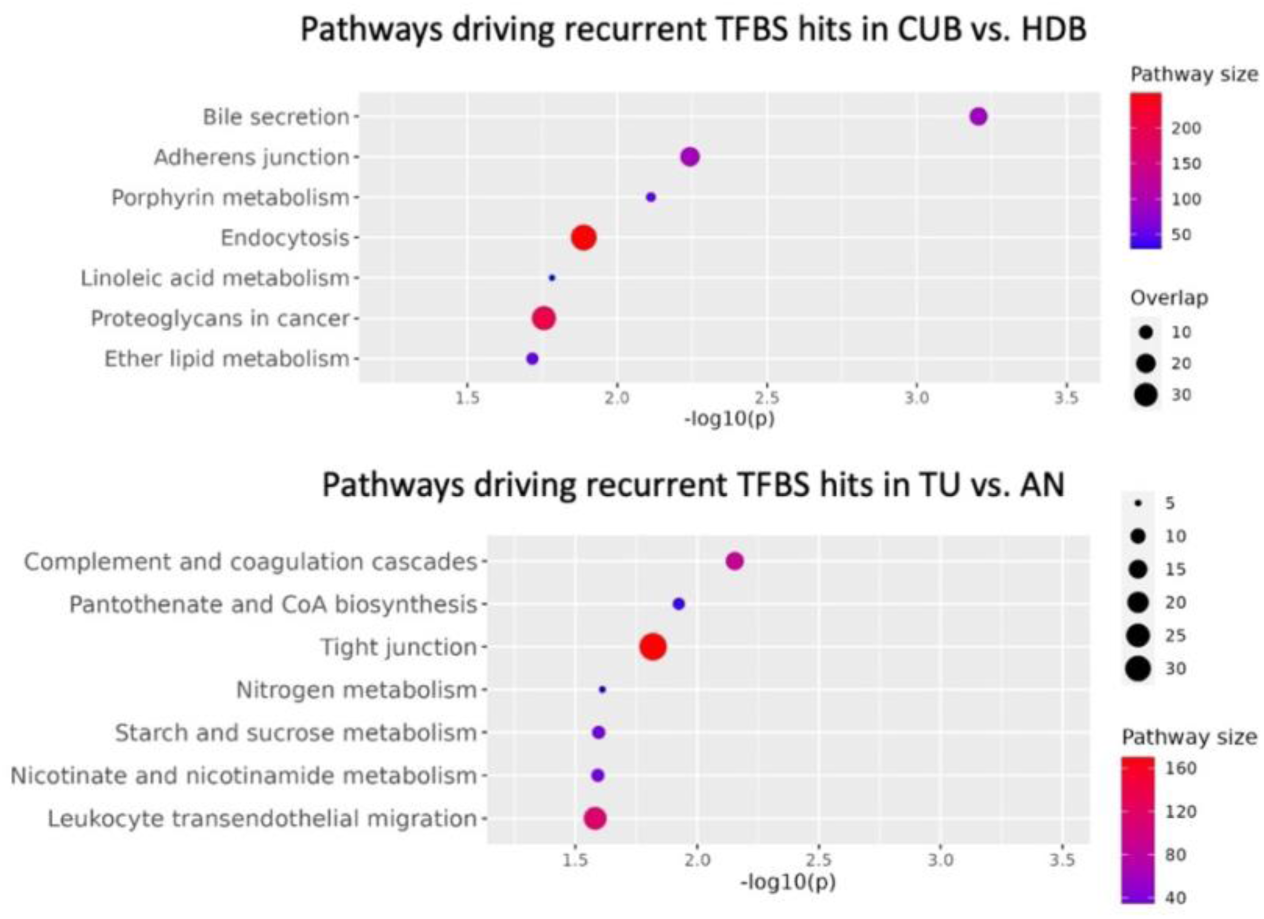
Pathways driving TFBS enrichment hits that are overlapping between the hypomethylated CpGs in the CUB vs HDB comparison and the TU vs AN comparison.

**Supplementary Figure 5.**
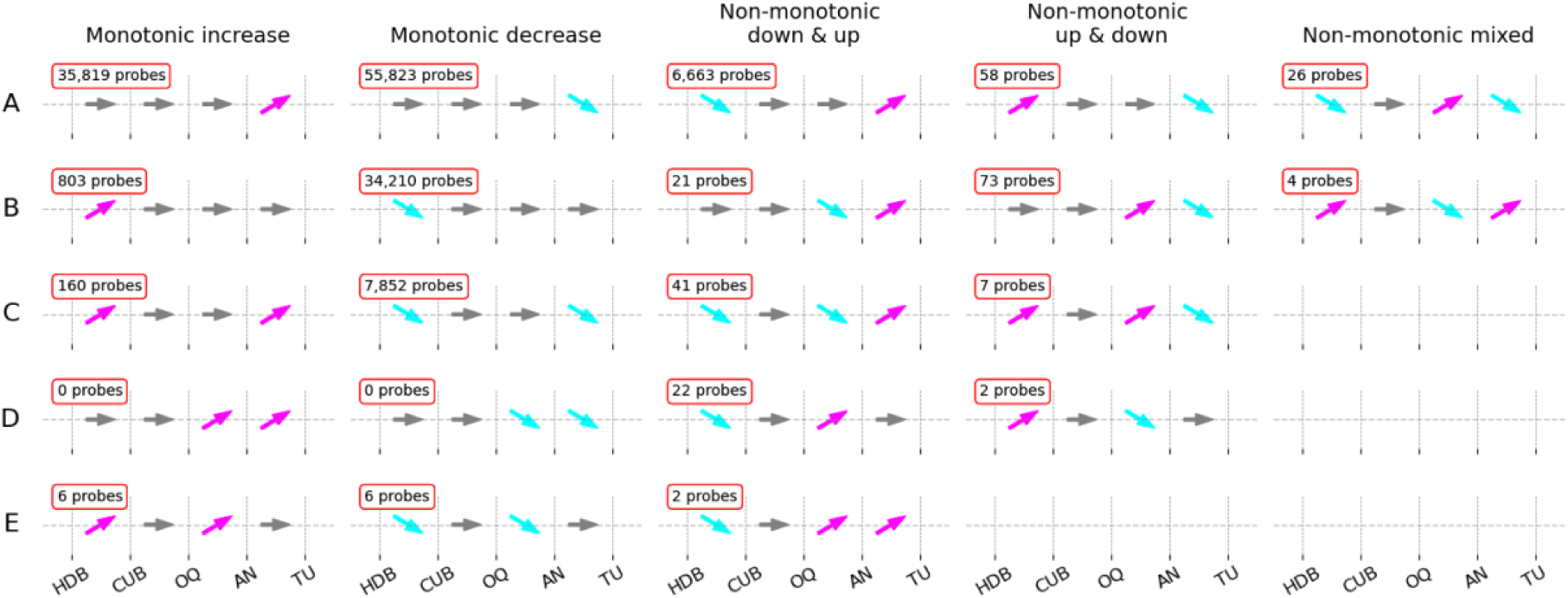
Differential methylation patterns along the TPxA and number of probes. The upward-pointing magenta arrows indicate hypermethylation, and the downward-pointing cyan arrows indicate hypomethylation. Each panel defines a single global trend along the TPxA, and red-lined text at the top left of each panel indicate how many probes fell into each global trend.

**Supplementary Figure 6.**
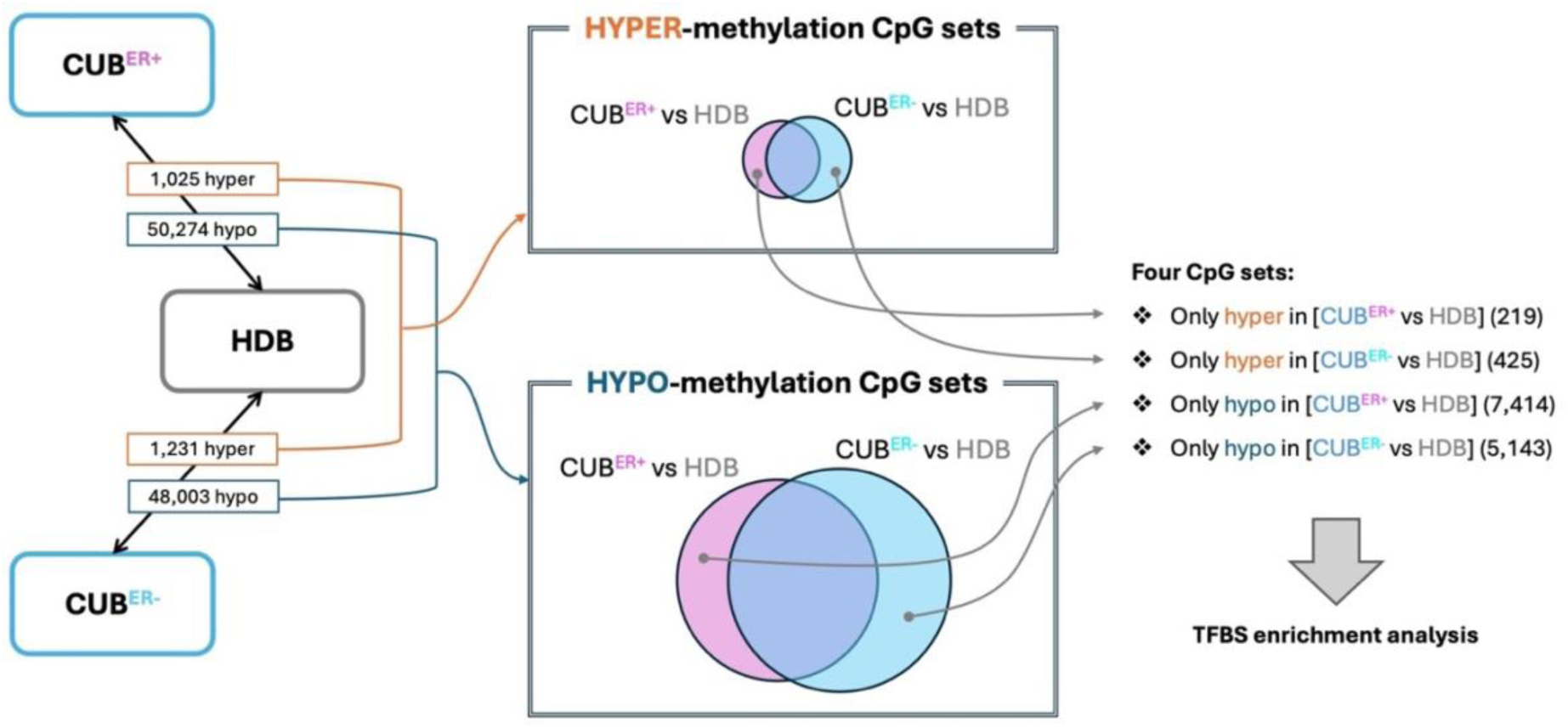
Analysis overview for CpGs that are uniquely DM in the CUB^ER+^ vs HDB or the CUB^ER-^ vs HDB comparison. HDB is always the reference category. Results of the TFBS enrichment analysis are shown in Supplementary Table 12.

**Supplementary Figure 7.**
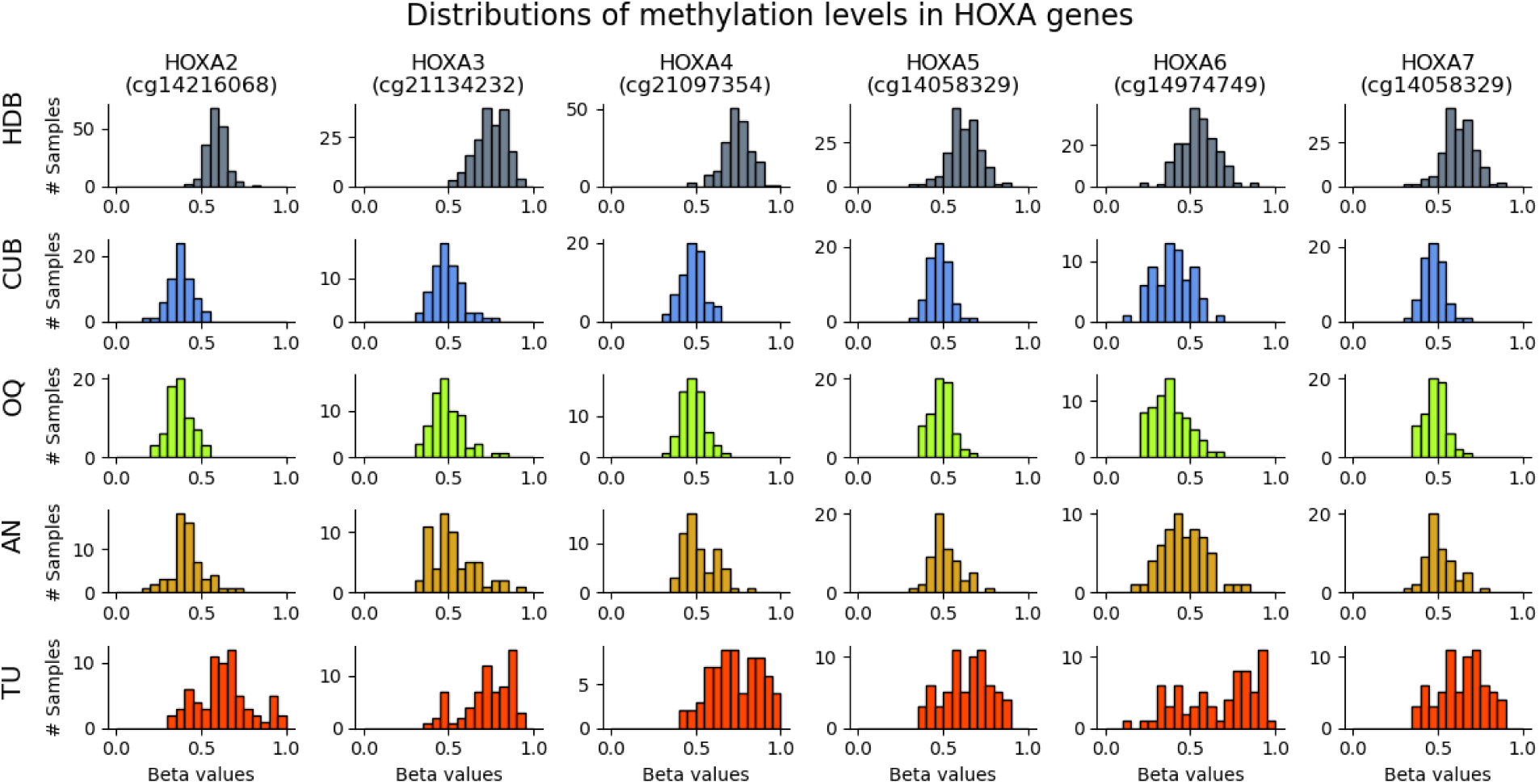
Beta value distributions of selected CpGs mapped to HOXA genes that were significantly hypomethylated in case-benign tissues.

## List of abbreviations

AN: Tumor-adjacent normal breast tissue
BMI: Body Mass Index
CpG: Sites in the genome where the sequence is cytosine followed by guanine
CUB: Contralateral unaffected breast tissue
DM: Differential methylation / differentially methylated
ER: Estrogen receptor
FDR: false discovery rate
HDB: Healthy donated breast tissue
HER2: Human epidermal growth factor receptor 2
HR: Hormone receptor
OQ: Opposite quadrant of the ipsilateral breast tissue
PCA: Principal component analysis
PRC2: Polycomb Repressive Complex 2
TF: Transcription factor
TFBS: Transcription factor binding site(s)
TPxA: Tumor Proximity Axis
tSNE: t-distributed stochastic neighbor embedding
TU: Tumor

## Declarations

### Ethics approval and consent to participate

This study was conducted with the approval of the Institutional Review Boards of Northwestern University (IRB #STU00051134) and UCLA (IRB #16-000853).

### Consent for publication

Not applicable.

### Availability of data and materials

The datasets generated and/or analysed during the current study will be available in NCBI’s Gene Expression Omnibus (GEO) under accession GSE287331 upon publication. (https://www.ncbi.nlm.nih.gov/geo/query/acc.cgi?acc=GSE287331)

### Competing interests

SAK serves as an Advisor for Havah Therapeutics. The remaining authors have no competing interests.

### Funding

This study was supported in part by the Bluhm Family Foundation, the Cancer Center Support Grant (CCSG) P30 CA060553, and the NIH grant R01LM013337.

### Authors’ contributions

SC and SAK conceptualized the study. TT conducted sample processing and experimental assays to generate data. MS and PG contributed data from a previous study. SD performed data processing, analyses, visualizations, and interpretations with guidance from SC, SAK, and WZ. SD drafted the manuscript, which was primarily edited by SC and SAK. All authors reviewed and approved the final manuscript.

## Acknowledgements

This research was supported in part through the computational resources and staff contributions provided for the Quest high performance computing facility at Northwestern University which is jointly supported by the Office of the Provost, the Office for Research, and Northwestern University Information Technology. We received support from the Pathology Core Facility of the Lurie Cancer Center which is supported by the Cancer Center Support Grant (CCSG) P30 CA060553.

